# Inhibition of platelet function by targeting the platelet cytoskeleton via the LIMK-signaling pathway

**DOI:** 10.1101/438051

**Authors:** Juliana Antonipillai, Sheena Rigby, Nicole Bassler, Karlheinz Peter, Ora Bernard

**Author notes:** Senior authors contributed equally to this work (Karlheinz Peter, Ora Bernard). Corresponding author: Juliana Antonipillai.

## Abstract

Actin is highly abundant in platelets, and platelet function is dependent on actin structures. Actin filaments are dynamic structures involved in many cellular processes including platelet shape changes and adhesion. The actin cytoskeleton is tightly regulated by actin-binding proteins, which include the members of the actin depolymerising factor (ADF)/cofilin family. LIM kinase (LIMK) and slingshot phosphatase (SSH-1L) regulate actin dynamics by controlling the binding affinity of ADF/cofilin towards actin. We hypothesised that inhibition of LIMK activity may prevent the changes in platelet shape during their activation and therefore their function by controlling the dynamics of Factin. Therefore, inhibition of LIMK activity may represent an attractive new strategy to control and inhibit platelet function; particularly the formation of stable platelet aggregates and thus stable thrombi.

## Introduction

Changes in platelet shape and functions are regulated by cytoskeletal rearrangements involving dynamic actin polymerisation and depolymerisation (1, 2). Platelets circulate in the blood as discs in an inactive form with a distinct organisation of microtubules (MTs) and actin filaments (F-actin). Actin comprises ~20-30% of the total platelet proteins (3). During activation, platelets undergo dramatic changes including a rapid shape change from a disc to a flat sphere; formation of pseudopods; disassembly of the peripheral microtubules ring; migration of the dense bodies towards the centre of the cell; secretion from its granules; conformational change and cytoskeletal anchorage of the integrin receptor αIIbβ3 (GPIIb/IIIa, CD41/61) and finally aggregation. All of these changes require alterations in the membrane skeleton as well as actin and tubulin remodelling (2).

The changes in the actin cytoskeleton are important for platelet activity in many cardiovascular functions. The dynamic process of the actin cytoskeleton is regulated by actin regulatory proteins including members of the actin depolymerising factors ADF/Cofilin. The activity of cofilin is regulated by the members of the LIM kinase family, LIMK1 & 2, that phosphorylate cofilin on serine 3 resulting in its inactivation (4, 5). LIMK1 and LIMK2 are serine protein kinases that are activated by phosphorylation of threonine 508/505, respectively, by the effectors protein kinases belonging to the Rho GTPase family that includes Rho kinase (ROCK) and p21 activating kinases (PAK 1 and 4) (6–8). Activated LIMKs phosphorylate and inactivate their substrate cofilin resulting in the accumulation of actin filaments within the cells (4, 5). On the other hand, the cofilin phosphatase slingshot (SSH1L) (also known as LIMK1 phosphatase) reactivates cofilin by dephosphorylation of both cofilin and LIMK1 (9, 10).

LIMK1 and SSH-1L play a major role in actin dynamics via the regulation of ADF/cofilin. Figure 1 represents a schematic diagram of actin filament formation via the downstream of ROCK/LIMK1 signal transduction pathway and the cofactors involved in the process.

**Figure 1:**
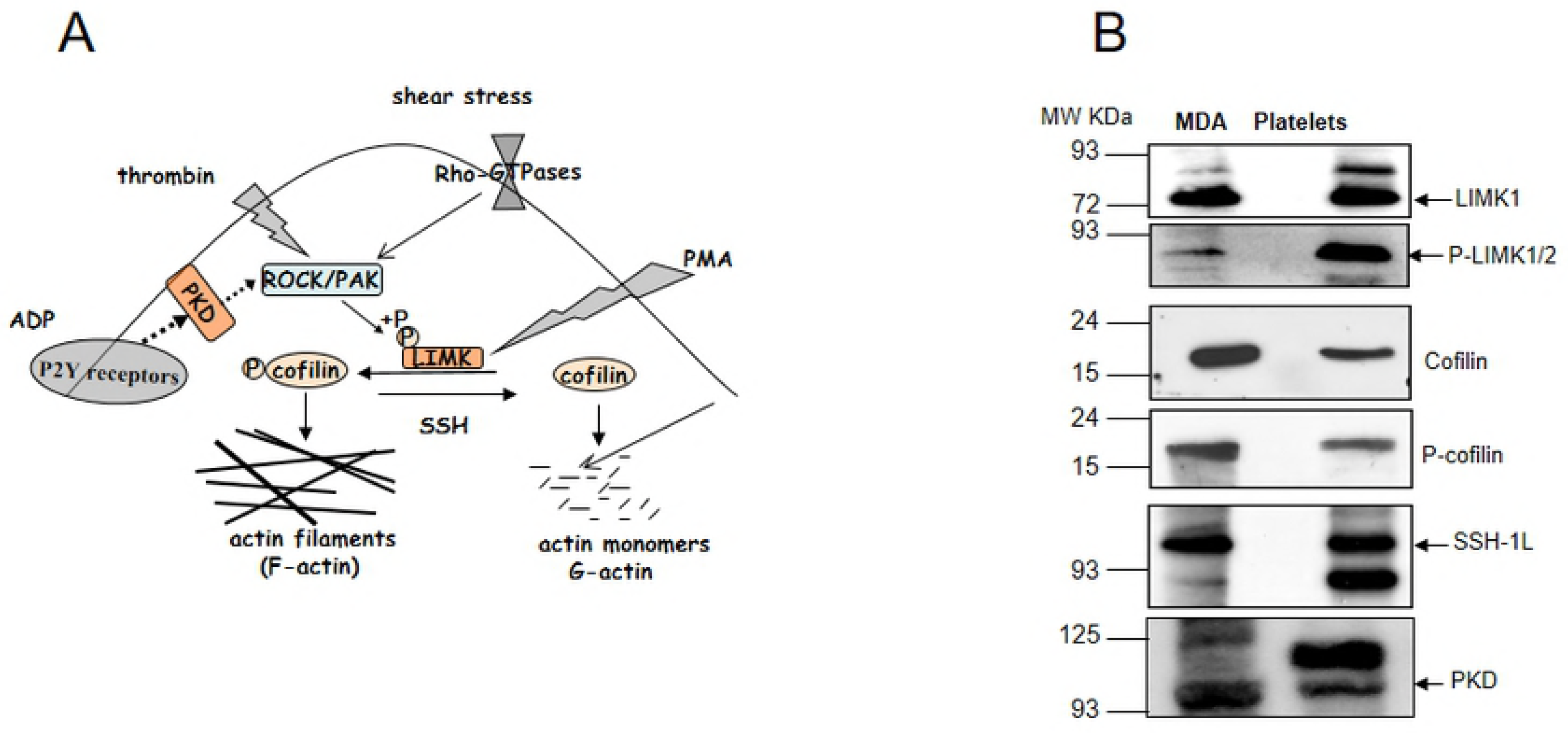
Schematic diagram of LIMK1 signal transduction and the expression of LIMK-associated proteins in human platelets. (A) LIMK1 promotes actin polymerization via inhibition of the actin depolymerizing factor, cofilin. Rho-associated kinase (ROCK) and p21-activated kinase1 and 4 (PAK1/4) phosphorylate and activate LIMK1, while slingshot phosphatase (SSH-1L) negatively regulates its activity. Some other possible LIMK1 activators are thrombin, ADP and PMA. (B) Platelets express LIMK1, cofilin, PKD and SSH-1L. Western blot analysis of purified platelets lysates probed with rat anti-LIMK1, rabbit anti-cofilin, anti-phospho-cofilin, anti-phospho-LIMK, anti-PKD and anti-SSH-1L. MDA-MB-231 human breast cancer cell lysate was used as positive control.

The finding by Pandey et al. that LIMK1 activity is important for thrombin-induced change of platelet shape and aggregation suggests that LIMK1 may be a new therapeutic target to inhibit platelet aggregation (11). Furthermore, these studies also demonstrated that ROCK activated LIMK1 during platelet shape change and aggregation. Interestingly, although LIMK1 was activated, no change in the level of P-cofilin was observed during shape change, while cofilin de-phosphorylation was observed during platelets aggregation (11). Other studies have demonstrated a role for thrombin in ROCK and LIMK1 activation. Addition of thrombin to human vein umbilical endothelial cells (HUVECs) resulted in ROCK activation followed by LIMK1 activation, increased P-cofilin and F-actin levels (12). Interestingly Aslan et al. demonstrated that also PAK, the other positive regulator of LIMK, regulates platelet shape change and aggregation (13).

The studies conducted by Estevez et al. reported that LIMK1 is involved in arterial thrombosis and that LIMK1 KO mice show no sign of bleeding complications (14). However, recent studies conducted by Kauskot et el. found that dysregulation of LIMK activity may be linked to macrothrombocytopenia (lower platelet counts) leading to bleeding complication in patients suffering from severe von Willebrand disease type 2B (15).

Here we demonstrate, that inhibition of LIMK by the small LIMK inhibitors, BMS3 (16) (17), Pyr1 (18) and 22j (19), can control platelet function.

## Materials and Methods

### Blood collection and platelet isolation

Platelet rich plasma (PRP) from human blood collected by venipuncture from healthy volunteers taking no medications and anti-coagulants was prepared as previously described. Washed platelets prepared from citrated PRP (1 ml) were passed through a pre-washed sepharose CL-2B column (Sigma) and eluted by addition of 1 ml of modified Tyrode’s buffer (150 mM NaCl, 2.5 mM KCl, 12 mM NaHCO_3_, 2 mM MgCl_2_, 2 mM CaCl_2_, 1mg/ml BSA, 1mg/ml dextrose; pH 7.4). After elution, platelets were diluted with modified Tyrode’s buffer either 1:50 for flow cytometry analysis or 1:5 for staining and adhesion assay purposes. For western blotting, PRP was acidified to pH 6.5 by addition of ACD buffer (1.32% w/v sodium citrate, 0.48% w/v citric acid, and 1.47% w/v dextrose) followed by addition of 10 mM Theophylline, to inhibit platelet activation during centrifugation. Acidified PRP was centrifuged at 720g for 10 minutes at 22°C, and the platelet pellet was washed with 5 mL of PBS containing 1 mM of Theophylline. After centrifugation, platelets’ pellet was resuspended in PBS containing Ca^2+^/Mg^2+^.

### Westernblots

Purified human platelets were lysed and analysed by immunoblot. Purified platelets also treated with ROCK or LIMK inhibitors for 5 minutes prior to immunoblot analysis. Membranes were probed with the following antibodies: rabbit anti-P-cofilin (cell signaling), rabbit anti-cofilin (cell signaling), rat anti-LIMK1 (20), rabbit anti-P-LIMK (Abcam), rabbit anti-P-MLC (cell signaling), rabbit anti-SSH-1L (kindly donated by Dr Bamburg JR) (10) and rabbit anti-PKD (Cell signaling). Samples were normalised against anti-actin or anti-GAPDH (Cell signaling).

### Static adhesion and immunostaining assays

Human platelets purified through Sepharose CL-2B column were added onto fibrinogen-coated cover slips blocked with 1% BSA for 1 hour at RT. Platelets, pre-incubated with the ROCK inhibitor (Y-27632), or LIMK inhibitor (22j) for 1 hour at RT, were allowed to adhere in the presence of 2 μM ADP at 37°C for 30 minutes. Coverslips were washed twice with Tyrode’s buffer and fixed with 1x Cellfix for 30 minutes followed by two washes with Tyrode’s buffer before mounting with Vectashield and visualized by microscopy (Nikon A1R confocal). Washed mouse platelets (1×10^7^/L in PBS) were added onto fibrinogen-coated cover slips as described above and permeabilized with 0.1% Triton X-100 for 30 minutes at RT followed by staining with Alexa^488^ phalloidin for 30 minutes at RT. Platelets were visualized using a Nikon A1R confocal microscope (Japan) Apo ×40 water 1.1 n.a objective. Stress fibres (F-actin) were visualized under 2D mode N-SIM structured illumination microscopy using a Nikon Ti inverted microscope (Japan) with an Apo ×100 oil 1.49n.a. objective and EMCCD camera (Andor Technology Ixon 3). The images were reconstructed using the NIS ELEMENTS software 4.10.

### Flow cytometry

Platelet rich plasma (PRP) isolated by gradient centrifugation from healthy human blood was incubated with and without ROCK inhibitor (Y27632, 20-300 μM) or LIMK inhibitor 22j (0-200 μM) for 5-10 min at 37ºC and then incubated with 2 or 20 μM ADP for 15 min at 37ºC. The platelets were stained with PE-conjugated mouse anti-CD62P mAb (P-selectin, anti-CD62P, clone AC1.2, BD Bioscience; 1:100) or PE-conjugated activation specific GPIIb/IIIa single-chain antibody (21) or FITC-mouse anti-PAC1 mAb (BD Bioscience; 1:100) for 15 minutes. Fixed platelets were analysed by a fluorescence-activated cell sorter (FACS Calibur, BD). The mean fluorescence indices (MFI) were analysed by CellQuest software.

### Platelet aggregometry

PRP aliquots of human blood pre-incubated for 1 hour with the ROCK or LIMK inhibitors were subjected to platelet aggregometry using an aggregometer (AggRam TM System) in the presence of 1 μM ADP.

### Clot retraction assay

PRP was incubated with 22j or solvent for 60 minutes at RT. Clot formation was initiated in glass tubes by the addition of 1 U/mL thrombin in the presence of 10 mM CaCl_2_. The retraction was measured after 1 hour. The extent of clot retraction was expressed as the volume of serum extruded from the clot as a percentage of the total reaction volume.

### F-actin/G-actin ratio assay

Washed platelets, treated with LIMK inhibitors, were lysed in actin stabilisation buffer (0.1 M PIPES, pH 6.9, 30% glycerol, 5% DMSO, 1 mM MgSO4, 1 mM EGTA, 1% Triton X-100, 1 mM ATP and protease inhibitors) on ice for 10 minutes. Samples were centrifuged in an ultracentrifuge at 100,000g for 1 hour, and the supernatant containing G-actin was recovered. The pellets containing F-actin were solubilized with actin depolymerisation buffer (0.1 M PIPES, pH 6.9, 1 mM MgSO4, 10 mM CaCl_2_ and 5 μM cytochalasin D). Aliquots containing the supernatant and pellet fractions were analysed by immunoblotting with HRP-conjugated anti-beta actin (Santa Cruz Biotechnology).

### In vitro kinase assay

The *in vitro* kinase assay was performed as previously described. Briefly, target proteins were purified and then subjected to an *in vitro* kinase assay for 30 minutes at 37°C in the presence of 5 μCi [^32^P] γ-ATP.

### In vitro phosphatase assays

Washed human platelets re-suspended in PBS were incubated with ADP (2 μM), Y27632 (20 μM) for 5 minutes at room temperature. Platelets were lysed in PBS lysis buffer [50 mM Tris-HCl (pH 7.4), 150 mM NaCl, 1% Triton X-100, Protease inhibitor cocktail] as described above. SSH-1L was immunoprecipitated with rabbit anti-SSH-1L polyclonal antibodies. Bacterially expressed purified GST-cofilin bound to glutathione sepharose beads was phosphorylated by *in vitro* kinase assay using active LIMK1 (Upstate) ^10^. Phosphatase assays were carried out at 30°C for 2 hours using 15 μL of ^32^P-GST-cofilin as a substrate for SSH-1L in a phosphatase buffer [30 mM Tris-HCl (pH 7.4), 30 mM KCl, 1 mM EDTA, 0.1 mg/ml BSA and protease inhibitors]. After centrifugation at 12,000g for 30 seconds, the radioactivity released by SSH-1L from ^32^P-GST-Cofilin was determined by β counter (PerkinElmer). The beads containing GST-^32^P-cofilin were subjected to western blotting. ^32^P-GST-cofilin was detected by a Phosphorimager and quantified using Image Quant probe software. The level of total GST-cofilin protein was determined by immunoblotting and quantified using the above software.

### Tail bleeding time

Mice were anaesthetized with ketamine (100 mg/kg), and xylazine (5 mg/kg) by intraperitoneal injection and the tip of their tail (5 mm) was cut and immediately immersed into saline, pre-warmed at 37°C. Bleeding time was monitored and recorded as the time needed for the cessation of the visible blood stream, for at least 1 minute. Maximum bleeding time was defined as 20 minutes and mice bleeding for more than 20 minutes were sacrificed. For drug administration, the right side jugular vein was bluntly exposed and catheterized. LIMK inhibitor 22j (19) (100 mg/kg), ROCK inhibitor Y-27632 (30 mg/kg, Sigma-Aldrich) or an equal amount of solvent or PBS were injected 1 minute before the tail was cut.

### Carotid artery thrombosis

Ferric chloride-induced injury in the carotid artery of mice was used as previously described (22). Mice were anaesthetized with ketamine (100 mg/kg) and xylazine (5 mg/kg) by intraperitoneal injection. Anaesthetized mice were placed under a dissecting microscope in a supine position, and a midline incision of the skin extending from the mandible to the sternal notch was made. The fascia was bluntly dissected up to the left common carotid artery. Thrombosis was induced by applying a piece of filter paper (1×3 mm, GB003, Schleicher & Schuell) saturated with 10% ferric chloride (Sigma) under the left common carotid artery for 3 minutes. A piece of plastic sheet (3×6 mm) was laid under the filter paper to prevent the absorption of ferric chloride by surrounding tissue. After removing the filter paper, the artery was rinsed with saline, and a nano-Doppler flow probe (0.5 VB or 0.5 PBS, Transonic) was positioned over the artery. The blood flow was measured by a flow meter (T106 Transonic) and recorded with a PowerLab data acquisition unit (AD Instruments). Thrombotic occlusion was defined by a drop in blood flow to 0.03 mL/min for at least 20 seconds, and reopening was defined by an increase in blood flow to 0.04 mL/min for at least 20 seconds. Maximum occlusion time was defined as 60 minutes and mice that had not occluded after 60 minutes were sacrificed.

### Drug administration

The right jugular vein was bluntly exposed and catheterized. All injections were conducted via this catheter. 100 mg/kg 22j or an equal volume of solvent was injected at 1 minute before the start of the ferric chloride injury. For reopening experiments, 500 U/g BW Urokinase and 100 mg/kg BW 22j or an equal volume of solvent were injected immediately after blood flow ceased.

### Statistical analysis

A Mann Whitney *t*-test (mean ± standard deviation are presented) was conducted on the occlusion experiments if time measurements were in the predefined range. If they were not within this range, a Gehan-Breslow-Wilcoxon test (median is presented) was conducted. A Gehan-Breslow-Wilcoxon test was also conducted on all bleeding time experiments. A Chi-square test was conducted on the reopening experiment. Differences were considered to be significant at p < 0.05. Other results are shown as means ± SEM from ≥ 4 independent experiments and from various donors. Groups were compared by t-test and a value of p < 0.05 was considered significant.

## Results

### Expression of the cytoskeleton regulators in platelets

It was previously demonstrated that LIMK1 protein is expressed in platelets (11). Using rat anti-LIMK1 monoclonal antibodies (mAb) (20), we have confirmed LIMK1 expression in platelets by western blotting (Figures 1B). Furthermore, we have shown that adult human platelets also express a number of other molecules that play roles in downstream of LIMK signaling pathway including cofilin, phospho-cofilin (P-cofilin), PKD and slingshot-1L (SSH1L) proteins (Figure 1B). As these proteins are involved in the regulation of actin polymerization, it is highly suggestive that they are important for the regulation of platelet function as previously suggested for LIMK1 and cofilin (11).

### LIMK plays an important role in platelet cofilin phosphorylation

Since cofilin is a major substrate of LIMK, the level of P-cofilin correlates with LIMK activity. We therefore tested the effects of LIMK and ROCK inhibition on cofilin phosphorylation after platelet incubation with the LIMK inhibitors 22j (19), BMS3 (17) (16) and Pyr1 (18) as well as the ROCK inhibitor Y-27632 (11) for 5 minutes (Figure 2). Incubation of platelets with these inhibitors resulted in a significant reduction in the level of P-cofilin, with 22j having the greatest effect on the levels of both P-cofilin and P-LIMK (Figure 2C). As the high concentration of 22j (Figure 2B) may also inhibit ROCK activity (19), we tested the effect of 10 and 100 μM 22j on the phosphorylation of the ROCK substrate, myosin light chain (MLC) using an anti-phospho-MLC (P-MLC) antibody. However, high concentrations of 22j did not affect MLC phosphorylation whereas it significantly reduced P-cofilin levels (Figure 2D), suggesting that the inhibition of LIMK activity by 22j is specific under these conditions.

**Figure 2:**
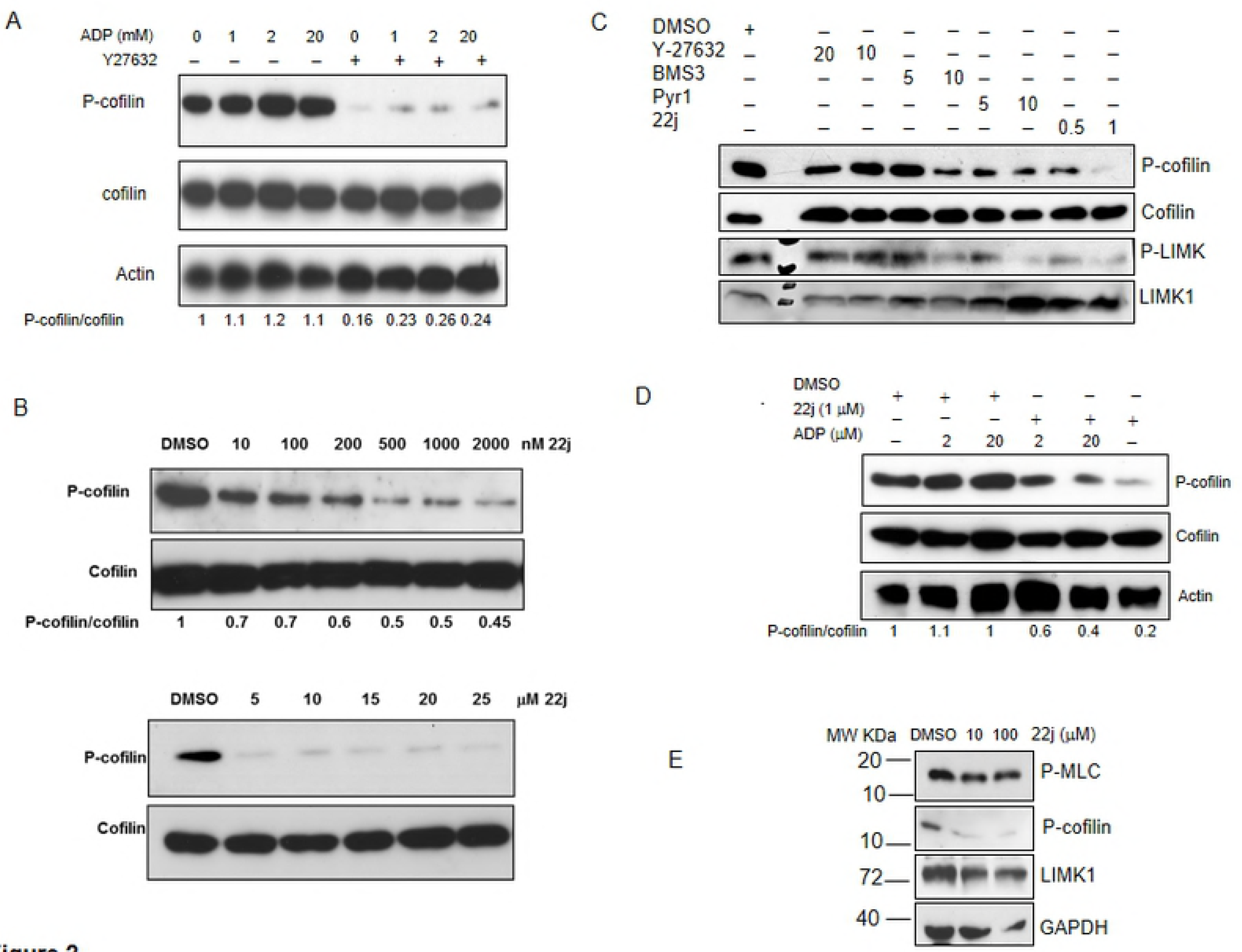
Pharmacological inhibition of LIMK activity results in inhibition of cofilin phosphorylation. (A) Inhibition of ROCK activity (direct upstream of LIMK) greatly reduced P-cofilin levels. Purified platelets were stimulated for 1 minute with ADP (0, 1, 2, 20 μM) or pre-incubated with 20 μM ROCK inhibitor Y27632 for 5 minutes prior to the incubation with ADP. Platelet lysates were immunoblotted with anti-P-cofilin Abs followed by probing the stripped membrane with rabbit anti-cofilin Abs. The numbers below the blots represent the ratio between the levels of P-cofilin to cofilin. (B) Purified platelets were incubated for 5 minutes with different concentrations of the LIMK inhibitor 22j (0-25 μM) and their lysates analyzed by immunoblotting with anti-P-cofilin and anti-cofilin antibodies. (C) Incubation of human platelets with LIMK and ROCK inhibitors greatly reduced P-cofilin and P-LIMK levels. Purified platelets were incubated for 5 minutes with two different concentrations of the LIMK inhibitors (BMS3, Pyr1 and 22j) and the ROCK inhibitor Y-27632 and their lysates analyzed by immunoblotting with anti-P-cofilin, anti-cofilin, and anti-P-LIMK antibodies. (D) Inhibition of LIMK activity greatly reduced P-cofilin levels. Purified platelets were stimulated for 1 minute with ADP (0, 2, 20 μM) or pre-incubated with 22j for 5 minutes prior to incubation with ADP. Platelet lysates were immunoblotted with anti-P-cofilin Abs followed by probing the stripped membrane with rabbit anti-cofilin Abs. The numbers below the blots represent the ratio between the levels of P-cofilin to cofilin. (E) The LIMK inhibitor 22j inhibits LIMK activity but not that of ROCK. Immunoblots of lysates prepared from platelets incubated for 10 minutes with 22j or DMSO were probed for phospho-MLC (P-MLC), P-cofilin and LIMK1. GAPDH was used as loading control.

### SSH-1L is responsible in part for the down regulation of P-cofilin levels after LIMK1 inhibition

We have demonstrated that inhibition of ROCK and LIMK1 activity by pre-treatment of platelets with the ROCK inhibitors as well as LIMK inhibitors for 5 minutes results in dramatic reduction in the level of P-cofilin, suggesting the involvement of a cofilin phosphatase in this process. We therefore compared the activity of SSH-1L, an established cofilin and LIMK1 phosphatase, in lysates prepared from untreated platelets and platelets treated with the ROCK or LIMK inhibitors. First, we confirmed that indeed the level of P-cofilin is reduced after treatment with these inhibitors and that SSH-1L is expressed in all samples (Figure 3A & C). SSH-1L activity was measured by the amount of ^32^P released from ^32^P-GST-cofilin by immunoprecipitated SSH-1L. We showed that the amount of ^32^P released from ^32^P-cofilin by SSH1L immunoprecipitated from the ROCK or LIMK inhibitor-treated platelets is greater than the amount released by SSH-1L purified from control platelets (Figure 3B & D).

**Figure 3:**
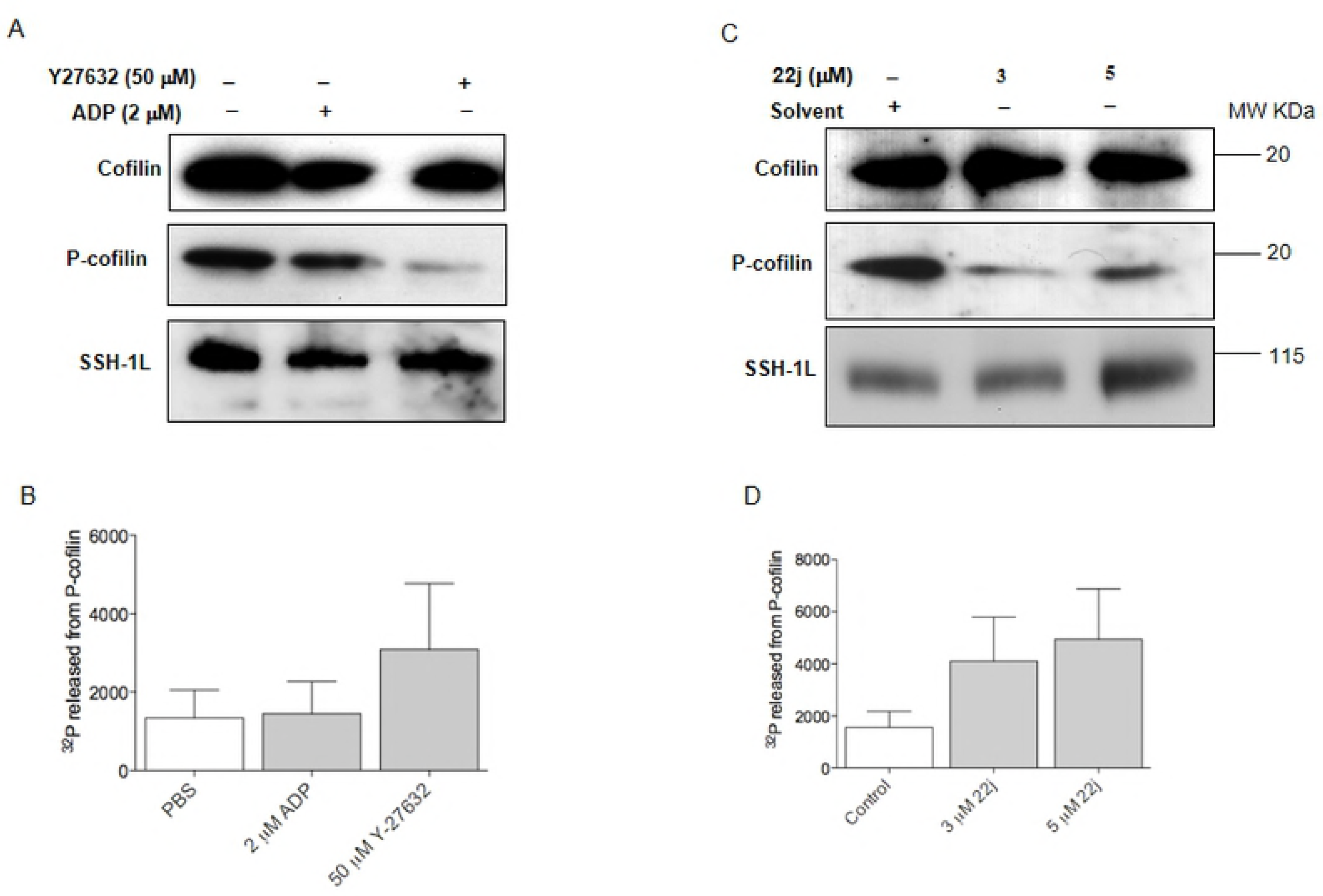
SSH-1L is responsible in part for the down regulation of P-cofilin levels after LIMK1 inhibition. (A) Immunoblot analysis of purified platelets treated with ADP or Y27632 for 5 minutes. The membrane was probed with rabbit anti-P-cofilin, anti-cofilin and anti-SSH-1L Abs. (B) Phosphatase assay analysis of SSH-1L immunoprecipitated from purified platelets treated with ADP or Y27632. Purified platelets were lysed after treatment with ADP and Y27632 for 5 minutes and SSH-1L was immunoprecipitated with anti-SSH-1L Abs and subjected to *in vitro* phosphatase assay using labelled ^32^P-GST-cofilin as a substrate. The amount of free ^32^P released from the labelled substrate was measured by β counter. The experiment was repeated 4 times with similar results and the average value was calculated. Bars represent mean ± SEM of 4 independent experiments. (C) Western blot analysis of platelets treated with 22j and subjected to SSH1L phosphatase assay. The membrane was probed with rabbit anti-P-cofilin, anti-cofilin and anti-SSH-1L Abs. (D) Purified platelets were lysed after treatment with 22j for 5 minutes followed by SSH-1L immunoprecipitation with anti-SSH-1L Abs. Immune complexes were subjected to *in vitro* phosphatase assay using ^32^P-GST-Cofilin as a substrate. The reaction was stopped by addition of SDS-loading buffer and subjected to western blotting. The amount of free ^32^P released from the labelled substrate was measure by β counter. The experiment was repeated 4 times with similar results and the average value was calculated. Bars represent mean ± SEM of 4 independent experiments.

### The ROCK/LIMK1 signaling pathway regulates platelet activity

To confirm the previous findings that the ROCK signaling pathway is important for the regulation of platelet function, we measured the activity of ADP-stimulated platelets in the presence of different amounts of the ROCK inhibitor Y27632 using P-selectin expression, a marker for platelet activation. P-selectin expression induced by 2 μM ADP was gradually decreased with increasing amounts of Y27632 (0-300 μM) (Figure 4A). The inhibition of ROCK activity with Y27632 partially inhibits P-selectin expression.

**Figure 4:**
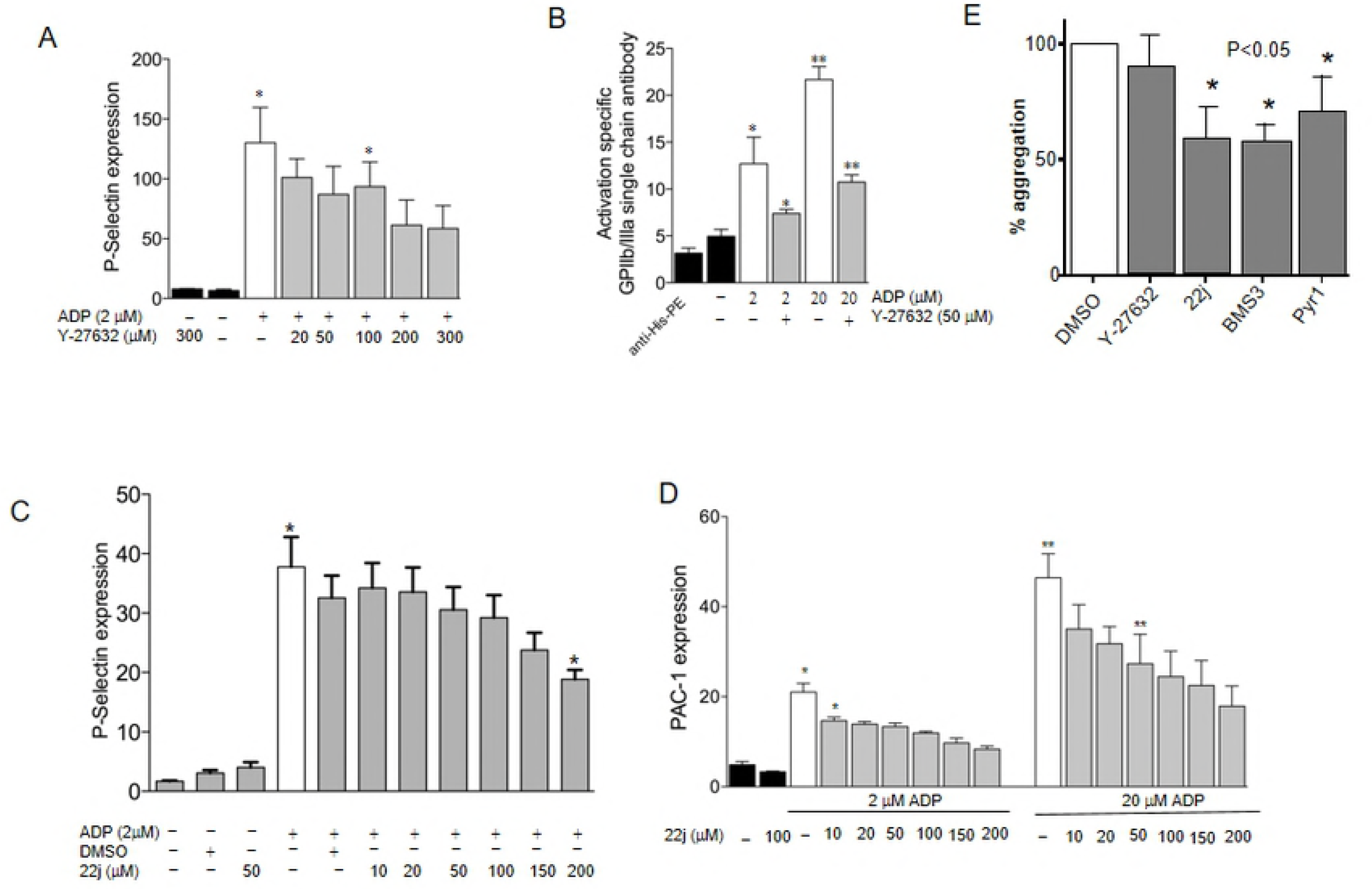
Platelet activation and aggregation are partially regulated via LIMK downstream signaling pathway. (A) P-selectin expression was measured during 2 μM ADP stimulation before and after incubation with different amounts of the ROCK inhibitor Y27632 (0-300 μM). Flow cytometry was performed following incubation with PE-conjugated mouse anti-CD62 mAb. These experiments were repeated more than 3 times, and the mean fluorescence intensity (MFI) was quantified. Bars represent mean ± SEM. Asterisks indicate P-value of <0.05. (B) GPIIb/IIIa activation during 2 or 20 μM ADP stimulation was measured using an activation-specific GPIIb/IIIa single chain antibody before and after incubation with Y27632. After incubation with the activation-specific GPIIb/IIIa single-chain antibody and subsequent incubation with anti-His-tag FITC-labelled secondary antibody, flow cytometry was performed. These experiments were repeated 3 times and mean fluorescence intensity (MFI) was quantified. Bars represent mean ± SEM. Asterisks indicate P-value of <0.05. (C) P-selectin expression was measured during 2 μM ADP stimulation before and after incubation with different amounts of the LIMK inhibitor 22j (0-200 μM). Flow cytometry was performed following incubation with PE-conjugated mouse anti-CD62 mAb. These experiments were repeated 3 times and mean fluorescence intensity (MFI) was quantified. Bars represent mean ± SEM. Asterisks indicate P-value of <0.05. (D) GPIIb/IIIa activation during 2 or 20 μM ADP stimulation was measured using a FITC-mouse anti-PAC1 mAb (BD Bioscience; 1:100) before and after incubation with 22j. These experiments were repeated 3 times and mean fluorescence intensity (MFI) was quantified. Bars represent mean ± SEM. Asterisks indicate P-value of <0.05. (E) LIMK is involved in platelet aggregation. Aggregation was measured in the presence of 2 μM ADP of PRP pre-treated with either LIMK inhibitors BMS3 (5 μM), Pyr1 (5 μM) and 22j (3 μM) or ROCK inhibitor Y-27632 (10 μM) or their solvent DMSO, for 1 hour at room temperature. The percentage of aggregated platelets after 10 minutes incubation: The plot shows data from 3 independent assays.

Another marker of platelet activation is the conformational change of the platelet integrin GPIIb/IIIa (23) (PAC-1). This receptor is constitutively expressed on platelets, however, upon platelet activation, it undergoes a conformational change that can be detected with a specific single-chain antibody. To further support our findings that ROCK activity is involved in platelet function, GPIIb/IIIa expression was measured by flow cytometry using His-tagged activation specific single-chain antibody (21). While ADP-stimulated human platelets result in GPIIb/IIIa activation, the addition of Y27632 decreases this level of activation by approximately 2 folds (Figure 4B).

### LIMK is involved in human platelet activity and aggregation

To confirm that the role of ROCK in platelet activity is via the regulation of the LIMK downstream signaling pathway, we investigated the changes in P-selectin and the conformational change of the platelet integrin GPIIb/IIIa (PAC-1) expression on ADP-stimulated platelets in the presence of different amounts of LIMK inhibitor 22j. Both P-selectin expression and GPIIb/IIIa (PAC-1) expression induced by 2 μM ADP was significantly decreased with increasing amounts of 22j (0-200 μM) (Figure 4C & 4D).

To further confirm our findings that the LIMK signaling pathway is important for platelet function, an aggregation assay was performed in the presence and absence of the specific LIMK inhibitors 22j (5 μM), BMS3 (5 μM), Pyr1 (5 μM) and the ROCK inhibitor Y-27632 (10 μM). Incubation with all these LIMK inhibitors led to a significant reduction in platelet aggregation (Figure 4E).

### Platelet adhesion is mediated by ROCK and LIMK

To test whether the ROCK/LIMK signaling pathway plays a role in integrin-mediated platelet adhesion, platelets pre-incubated with 20 μM Y-27632 or 3 μM 22j were activated with 2 µM ADP and allowed to spread on fibrinogen-coated cover slips. Incubation with Y-27632 and 22j reduced the number of platelets that adhered to fibrinogen (Figure 5A and B). These results suggest that platelet adhesion to fibrinogen occurs via activation of ROCK signaling cascade mainly via LIMK.

**Figure 5:**
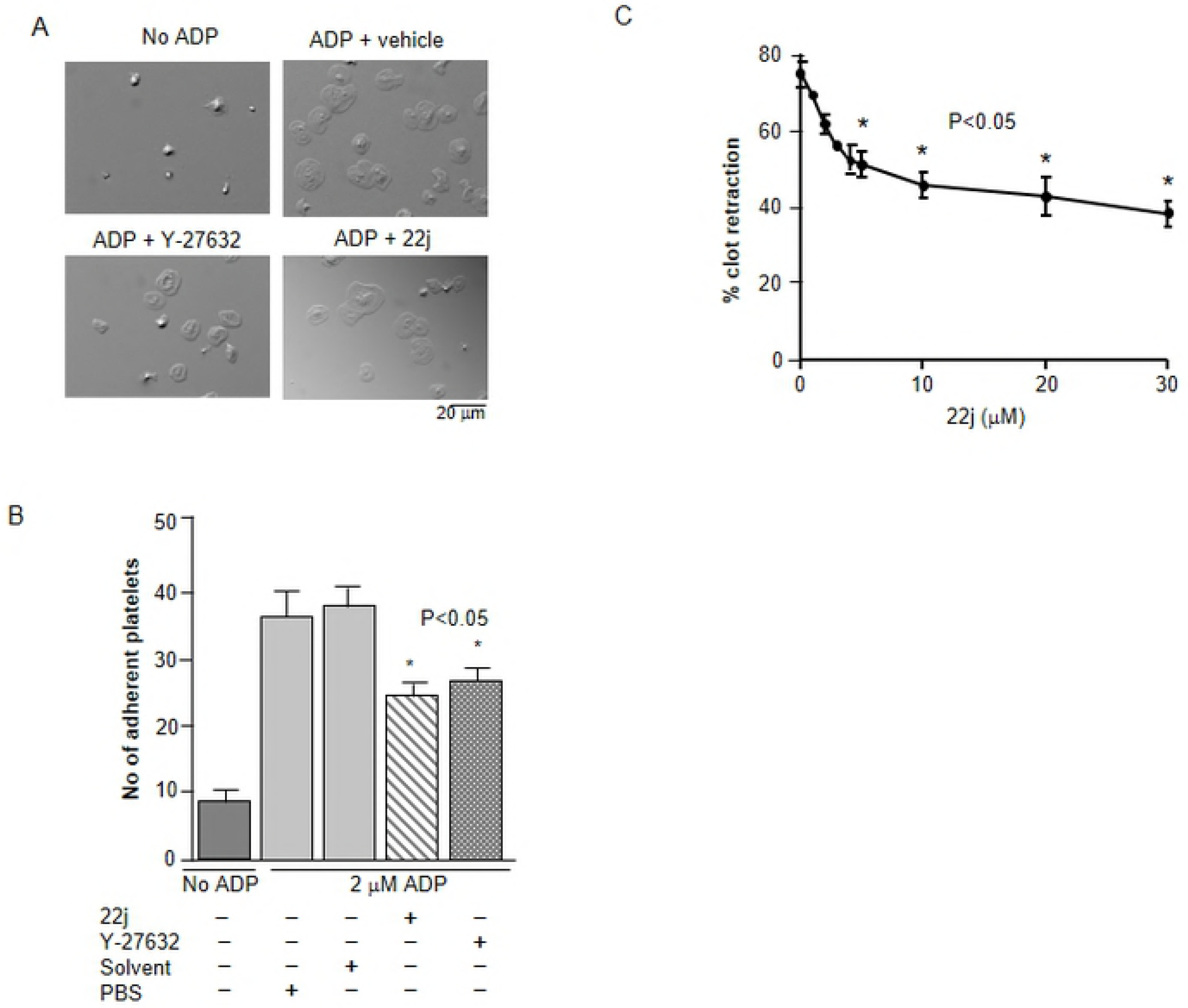
Platelet adhesion is inhibited by the ROCK and LIMK inhibitors. (A) Platelets pre-treated with Y-27632 or 22j for 1 hour were allowed to adhere onto fibrinogen-coated cover slips for 30 minutes in the presence of 2 μM ADP. Adherent platelets were visualized by light microscopy. The assay was repeated at least three times with similar observations. Bar=50 μm. (B) The average number of adherent platelets counted from 12 images generated from more than 3 independent experiments. Bars represent mean ± SEM, *P<0.05. (C) The LIMK inhibitor 22j reduces clot retraction. PRP were pre-treated with 22j for 1 hour at room temperature and subjected to clot retraction assay. The average volumes of serum extruded from the clot as the percentage of total reaction volume were calculated from triplicate samples.

### ROCK/LIMK control the actin cytoskeleton during platelet spreading

To investigate if LIMK is involved in regulating the tensile strength of the actin cytoskeleton, a clot retraction assay was performed with PRP in the presence of 22j. A significant reduction in platelet clot retraction was observed with increasing concentrations of the LIMK inhibitor 22j (Figure 5C). To confirm our findings that platelet actin polymerization is regulated by LIMK, F-actin was visualized using fluorescence microscopy. Platelets treated with 20 μM Y-27632 or 3 μM 22j displayed reduced adhesion (Figure 5A & 6A), reduced F-actin content (Figure 6B) and reduced spread surface area (Figure 6C). Stress fibre formation in platelets treated with either 22j or solvent was resolved with structured illumination microscopy (Figure 6D). Platelets treated with 22j showed an accumulation of chopped or truncated F-actin fragments at various lengths. However, we observed only a small trend towards reduction of the F-actin/G-actin ratio for both Pyr1 and 22j treated platelets compared to control platelets (Figure 6E and F).

**Figure 6:**
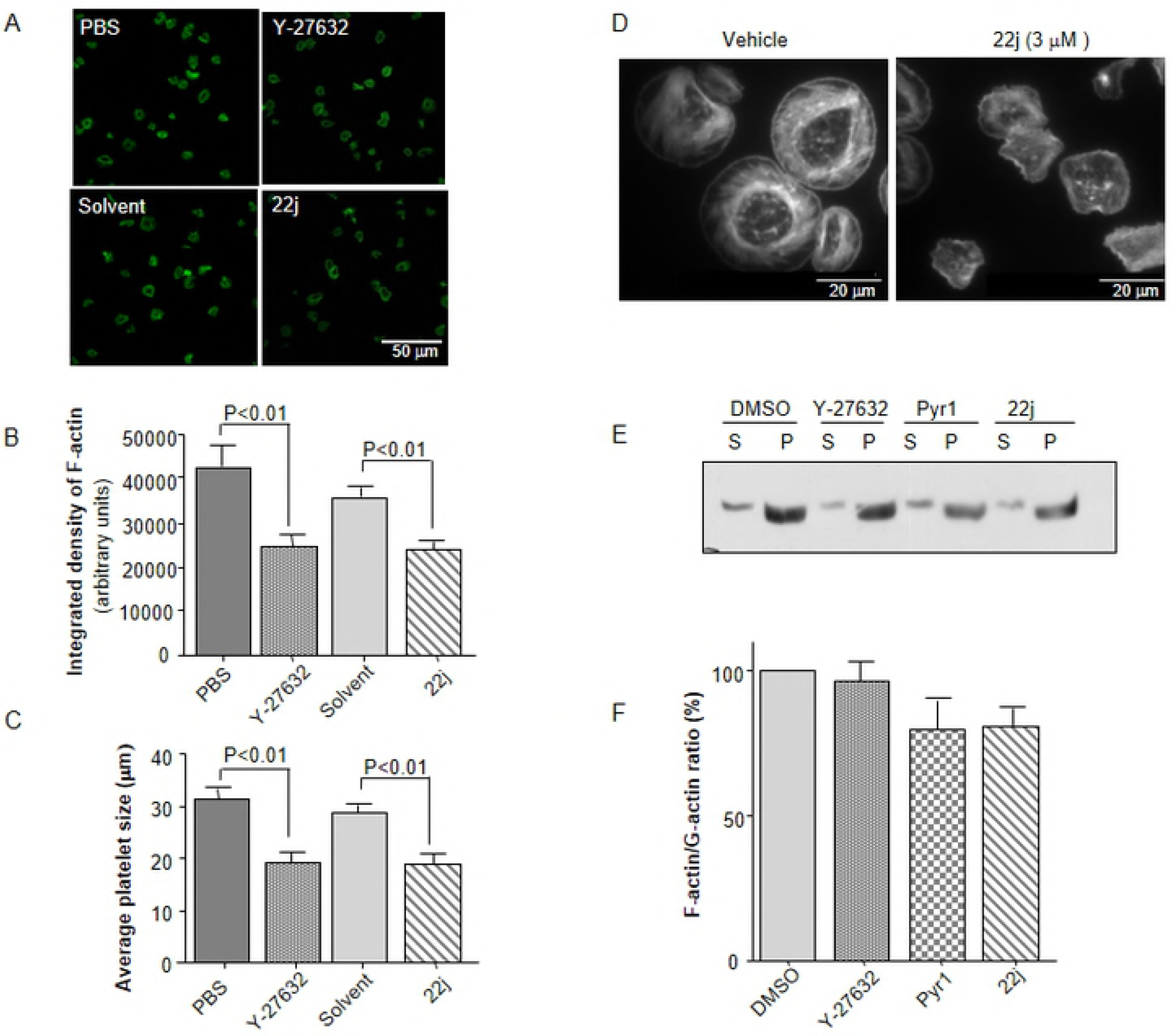
Stress fibres formation and platelet shape are regulated via downstream of ROCK/LIMK signaling. (A-C) Incubation with Y-27632 or 22j reduces platelet spreading and affects their shape. Platelets were pre-treated with Y-27632 or 22j or with vehicle for 1 hour then allowed to spread on fibrinogen-coated cover slips in the presence of 2 μM ADP. Adherent platelets were fixed, permeablized and then stained with Alexa^488^ phalloidin for F-actin. (A) Stained platelets were visualised under Nikon A1R fluorescence microscope. Bar=50 μm. (B) Incubation with Y-27632 and 22j reduces the surface area of spreading platelets. The average size of the platelets was calculated from more than 12 images generated from 3 independent experiments using Fiji software. (C) The integrated density representing F-actin density is reduced in Y-27632 or 22j treated platelets. The average integrated density of the adhered platelets was calculated by Fiji software. (D) Platelets spreading and stress fibres formation were affected by 22j. Alexa^488^ phalloidin stained platelets were visualized under 2 D mode Nikon N-SIM structured illumination microscope. Bar=20 μm. (E) Total F-actin/G-actin ratio was slightly reduced by LIMK inhibitors. Platelets were treated with Y-27632 or 22j or their vehicle before the standard F-actin/G-actin assays were performed. The samples were analyzed by Western blots probed with anti-actin-HRP antibodies. (F) The F-actin/G-actin ratios were calculated from 3 independent assays.

### Effects of LIMK inhibition on tail bleeding and thrombus occlusion time in mice

To further explore the involvement of LIMK in platelet function and its potential therapeutic implication, we performed tail-bleeding time assays with C57Bl/6 WT mice treated with 22j (100 mg/kg) or Y-27632 (30 mg/kg). A significantly prolonged bleeding time was observed in WT mice treated with 22j and Y-27632 (Figure 7A). Bleeding in the control mice stopped within 1-3 minutes (p< 0.01) while that of the Y-27623 and the 22j treated mice stopped after 15 and 20 minutes, respectively (Figure 7A). In addition, the carotid artery occlusion time following topical exposure to 10% ferric chloride in control mice was 662.0 ± 85.19 seconds, while the occlusion time of 22j administered mice was significantly increased to 998.6 ± 298.6 (p< 0.01) (Figure 7B). Similarly, the occlusion time of the Y-27632 administered mice was significantly increased from its PBS control 534 ± 59.14 to 1412 ± 284.2 seconds (p< 0.02). Our *in vitro* studies of pharmacological LIMK inhibition corroborate our *in vivo* experiments demonstrating that LIMK inhibition impairs platelet function resulting in a prolongation of bleeding and occlusion times.

**Figure 7:**
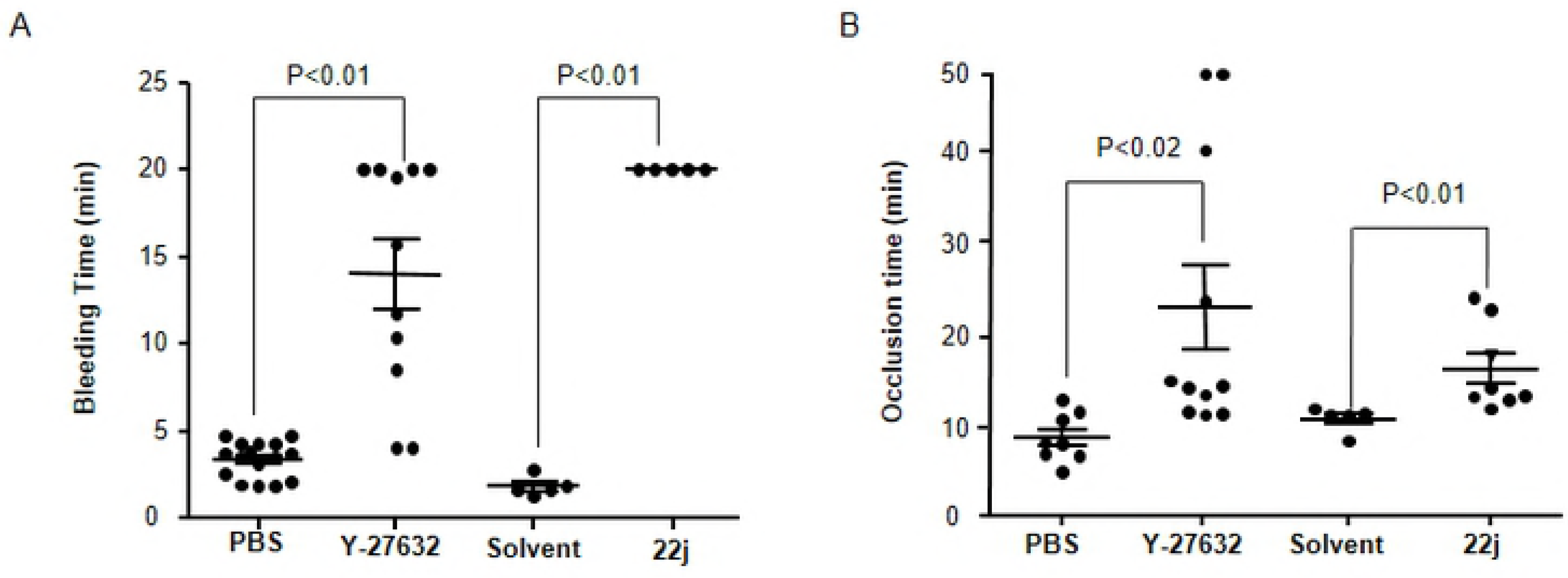
LIMK regulates platelet function and haemostasis in mice. Tail trans-section was performed at 5 mm from the tip of the tail and bleeding time was monitored. (A) The bleeding time of the Y-27632 and the 22j treated mice was significantly prolonged in comparison with that of control mice. (B) The carotid artery occlusion times in the 22j and Y-27632 treated mice were significantly longer than that of the control mice.

### 22j is a potential facilitator of thrombolysis

To investigate whether 22j can facilitate thrombolysis, we used ferric chloride to induce a thrombotic occlusion of the carotid artery in wild type mice followed by an injection of 500 U/g body weight (BW) urokinase, which is a clinically used fibrinolytic drug, and monitored the time taken for the vessel to reopen. When 22j (100 mg/kg) was injected simultaneously with urokinase, 50% of the mice showed reopening of the vessel compared to only 14% when injected with urokinase alone (n=14 each, p<0.05). This indicates that 22j may be a potential drug candidate for facilitating thrombolysis in combination with a fibrinolytic drug.

## Discussion

Anti-platelet therapy is highly beneficial for patients with cardiovascular diseases such as myocardial infarction and stroke. However, bleeding complications with current drug therapies can prevent their use in many patients that would otherwise benefit from them. The aim of this study is to understand the basic mechanisms involved in the regulation of the platelet cytoskeleton in order to identify new targets and novel strategies for anti-platelet therapy, which preferentially target thrombus stabilisation.

Actin and tubulin comprise the majority of the platelet proteins and changes in their polymerization state are essential for platelet shape change, granule secretion and aggregation. Inhibition of actin polymerization has been shown to reduce the stability of platelet aggregates (24). It is therefore likely that cytoskeletal regulators such as LIMK and its phosphatase SSH-1L are involved in platelet aggregation and play a role in the formation of stress fibres and filopodia, resulting in stabilization of platelet thrombi.

Pandey et al. demonstrated that during platelet aggregation the signal transduction downstream of ROCK plays a significant role in altering platelets function via the control of F-actin levels, in thrombin-activated platelets (11). These authors also demonstrated the involvement of LIMK1 in thrombin-stimulated platelets (11). In agreement with that study, we confirmed the expression of LIMK1 and cofilin in resting human platelets (Figure 1B).

We have further investigated the involvement of LIMK in the regulation of platelet function by using LIMK specific inhibitors. We observed reduced levels of LIMK and cofilin phosphorylation in platelets treated with the specific LIMK inhibitors BMS3 (17), Pyr1 (18) and 22j (19). We also noticed a rapid drop in P-cofilin levels with increasing amounts of 22j (Figure 2B). We have also observed that inhibition of LIMK activity by 22j or the ROCK inhibitor Y27632 control platelet adhesion as well as F-actin levels (Figure 5 & 6). Furthermore, inhibition of PAK activity by a PAK specific inhibitor was shown to control platelet spreading and F-actin levels (25). PAK and ROCK are two well-established upstream regulators of LIMK. Here we show that the inhibition of LIMK activity by the LIMK specific inhibitors inhibits platelet aggregation (Figure 4E), indicating that LIMK plays an important role in platelet function through control of F-actin levels, as previously demonstrated for thrombin-activated platelets by Pandey et al. (11).

During platelet activation by increasing concentrations of ADP, there was no change in the level of cofilin phosphorylation. However, when platelets were pre-incubated with the ROCK or LIMK inhibitors (Figure 2) cofilin phosphorylation was dramatically reduced. As mentioned above, the dramatic change in p-cofilin levels cannot be explained only by the reduction in LIMK1 activity. We therefore propose that the increased activity of cofilin phosphatase is responsible for the dephosphorylation of cofilin within 5 minutes inhibition of ROCK, or LIMK. As SSH1L is expressed in platelets, it is reasonable to suggest that it may play a role in the regulation of p-cofilin levels in platelets. Indeed, the experiments presented in Figure 3 indicate that SSH1L activity is increased after incubation with the ROCK inhibitor suggesting that inhibition of this enzyme results in SSH1L activation. We and others have previously demonstrated that SSH1L activity is regulated by phosphorylation (10). Eiseler et al. (26) have shown that phosphorylation of SSH1L by protein kinase D1 (PKD1) at Serine 978 generates a 14-3-3 binding site. SSH1L binding to 14-3-3 inhibits its interaction with F-actin. As SSH1L activity is greatly enhanced by its binding to F-actin, inhibition of this association results in down regulation of SSH1L activity (26). PKD1 is activated by the Rho-ROCK pathway, suggesting that inhibition of ROCK may result in PKD1 inhibition and up-regulation of SSH1L activity leading to cofilin dephosphorylation after ROCK inhibition. Indeed, we have shown that PKD is expressed in platelets (Figure 1B). We have previously demonstrated that LIMK1 interacts with SSH1L resulting in its dephosphorylation (10). It is possible that binding of LIMK1 to its inhibitor releases the interaction between these two enzymes favouring the interaction between SSH-1L and cofilin resulting in increased SSH-1L activity towards cofilin. Interestingly Leonard *et al.*, have demonstrated that both LIMK and SSH-1L have a selective role in endothelial cell inflammation associated with intravascular coagulation (27).

The studies conducted by Estevez et al. reported that LIMK1 is involved in arterial thrombosis, and LIMK1 KO mice show no sign of bleeding complications (14). In contrast, our studies with the LIMK inhibitor treatments did show significant bleeding prolongations. Interestingly, relatively high levels of P-cofilin protein in LIMK1 KO platelets were found by Estevez et al. (14) suggesting that either their LIMK1 KO model is not a complete knockout, and the kinase domain may expressed as previously suggested by Bernard (28), or LIMK2 is expressed in platelets, and that phosphorylates cofilin. However, previously Pandey et al. demonstrated that LIMK2 is not expressed in platelets (11). Till to date, LIMK1 and LIMK2 are the only known kinases that phosphorylate cofilin. We have been previously demonstrated the complete inhibition of phospho cofilin by knocking down of both LIMK 1, and LIMK2 with siRNA in human cell lines including breast cancer cells (16). The LIMK inhibitors that we used in this project, 22j, pyr1 and BMS3, inhibit both LIMK1 and LIMK2 and therefore, we can conclude that both LIMK1 and LIMK2 have a significant role in platelet function.

Therapeutic targeting of the platelet cytoskeleton is an attractive novel approach for anti-platelet therapy. Thrombus destabilization in the clinical setting of thrombolysis, for example in patients with myocardial infarction, is of particular interest. Whereas fibrinolytic drugs can dissolve the extracellular fibrin mesh, a LIMK inhibitor could destabilize the intracellular cytoskeleton. Indeed, we have shown that the LIMK inhibitor 22j in combination with urokinase was more efficacious than urokinase alone in restoring blood flow in the carotid artery following vessel injury and thrombotic occlusion. This opens up a new strategy in pharmacological thrombolysis that warrants to be tested further.

## Acknowledgements

We thank Dr Bryce Harrison (Lexicon) for his generous gift of 22j, Dr Laurence Lafanechere for her generous gift of Pyr 1, Prof James Bamburg for his generous gift of the anti-SSH-1L Abs and Dr Iska Carmichael (Monash Micro Imaging) for excellent technical assistance with fluorescence microscopy.

